# Nucleotides inclusion in pet food: effect of thermic treatment and shelf-life

**DOI:** 10.1101/353813

**Authors:** N. Russo, A. Candellone, D. Galaverna, L. Prola

## Abstract

A growing trend of nucleotides supplementation in pets is occurring in recent years with the final goal of reinforcing the immune system and promoting intestinal function. Despite that, data evaluating possible alterations of nucleotides during pet food processing are lacking. The aim of the study was to evaluate the recovery percentage of nucleotides in dry and canned food after processing and storage. Selected dry and canned feed were supplemented with 0.4/100 g dry matter basis of Prosol petMOD™. Recovery percentage in dry food was 75% and 74% at the end of technological process and a 12-month shelf-life, while in canned food it was 43% and 41%, respectively. These results support the hypothesis that N could be included during pet food process but industries have to increase the awareness about possible losses, especially in canned food.

## Introduction

In recent years there has been an increase in nucleotides supplementation, in particular in baby food and in parenteral preparations, as numerous researches have demonstrated that including nucleotides in such products lead to a reinforcement of the immune system and to a healthier gut in the receiver [1].

Nucleotides should be considered as “semi-essential” nutrients. It means that, in maintenance condition the endogenous production satisfy requirements while in puppies and in course of tissue damages, the exogenous administration is needed.

They are phosphoric nucleoside esters, formed of three components: a weakly basic nitrogenous compound, a pentose sugar and one or more phosphate groups. They constitute the basic units of the nucleic acids DNA and RNA [2–3]. Most important are adenosine, guanosine, inosine, cytidine and uridine monophosphates. Generally, there are three potential sources of nucleotides: de novo synthesis, salvage pathways and dietary nucleotides [4– 5].

De novo synthesis of nucleotides is a metabolically costly process requiring substantial amounts of energy in the form of ATP [2]. Another mechanism for maintaining the cellular nucleotide pools isthe salvage pathway. This pathway recycles 90% or more of the purine bases. Thus, it is suggested that the salvage pathway is dependent on the availability of free purine and pyrimidine bases [4]. This pathway requires less energy than the reactions needed for the de novo synthesis of nucleotides, and is characterized by linkage of a ribose phosphate moiety to free bases formed by hydrolytic degradation of nucleic acid and nucleotides. Furthermore, some tissues have limited capacity for the de novo synthesis of nucleotides, thus requiring exogenously supplied bases that can be utilized by the salvage pathway [6–7].

For example, the intestinal mucosa, hematopoietic cells of the bone marrow, leucocytes, erythrocytes and lymphocytes are incapable of de novo production (Sanderson & He, 1994), and thus utilize the salvage pathway, suggesting that an exogenous supply of nucleotides via diet might be important for aforementioned cells [4].

So far, different studies on nucletotides supplementation are available in farm animals, while few studies had been conducted in pets. Rutherfurd-Markvick et al. [9], demonstrated that 43 cats fed with an integrated diet with nucleotides showed an increased proliferative response of post-vaccination lymphocytes while other authors showed an improvement of the immune system in puppies fed with supplemented food [10–11] or in dogs suffering from leishmaniasis [12]. Given the above and in the view of a commercial growing trend of nucleotides supplementation also in pets, it is important to properly understand if and how nucleotides can undergo alterations during pet food processing and storage.

Pet food available on the market could be manufactured by extrusion/expansion (dry food) process and retort sterilization (canned food). The traditional dry pet food is obtained from the extrusion of a finely ground mixture of ingredients of animal and vegetable origin, in variable percentages depending on the species (dog or cat), age or size of the animal to which the food is destined. The extrusion process is fundamental to favor starch digestibility considering that cats and dogs have an incomplete enzyme pattern dedicated to this function [13]. To obtain this result, the food products, after addicting water and steam, undergo a short but intense heating and mechanical treatment. To guarantee storage at room-temperature, dry pet food is then dehydrated in stoves for 20-50 minutes. This industrial process, during which food components are exposed to high temperatures, could lead to a degradation of all heat-sensitive and easily oxidable nutrients. [14].

From a technological point of view, the canned pet food is considered a preserved one with a pH close to neutrality; to guarantee its microbiological, chemical and physical stability and a prolonged shelf -life, it has to be subjected to very intense thermal treatments. For this reason, over-dosages of some thermolable ingredients, such as nucleotides, are necessary to guarantee an adequate final recovery net of losses due to heating [15].

The aim of the present work was to study nucleotides inclusion in a dry and a canned pet food, determining their recovery percentage at the beginning and at the end of the technological production and after a 12-month shelf-life, in order to underline, whether or not, these components could experiment significant degradation after heating and prolonged storage.

## Material and Method

### >Preparation of dry food

A commercial dog kibble formulation was prepared with a percentage of inclusion of 0.14 g/100 g on DMB of Prosol petMOD ™, a *Kluyveromices fragilis* yeast cell derivative containing a high concentration of free nucleotides 5’-mono phosphate (> 40%) and total nucleic acids (> 80%), produced by PROSOL S.p.A. (Madone, 24040 BG - IT). Supplemented nucleotides were adenosine (AMP), cytidine (CMP), uridine (UMP) and guanosine (GMP) in a pool depicted in Table 1.

**Table 1.**
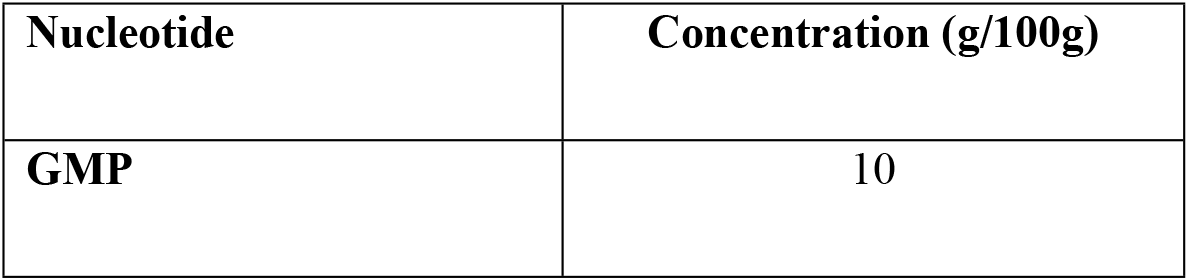
F.ree 5’-mono-phosphate nucleotide pool in Prosol petMOD ™.

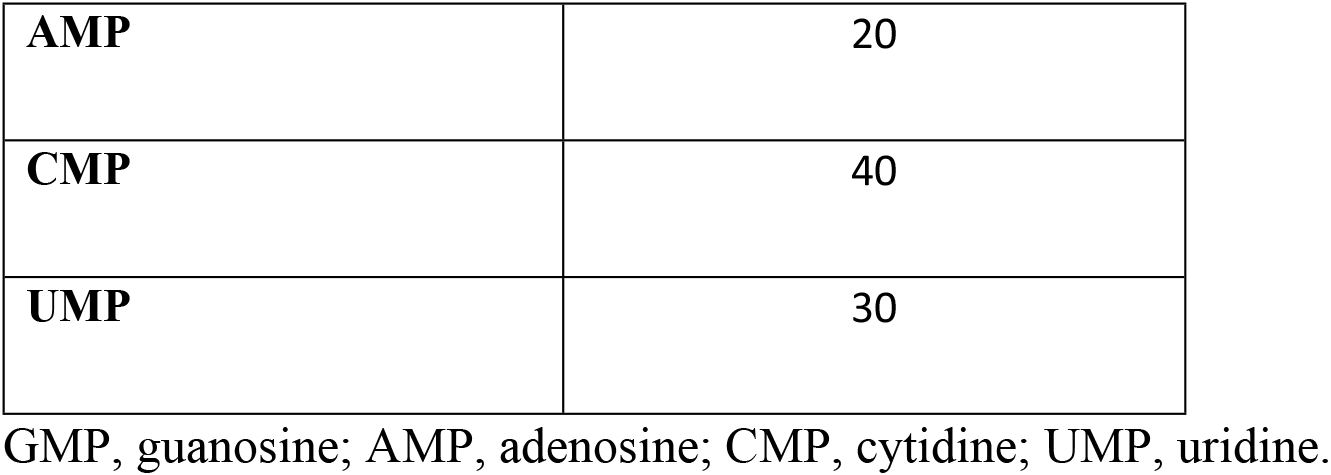

Table 2 summarizes the production flow occurred during dry food preparation. In detail, nucleotides inclusion occurred during the mixing phase. A customized dry pet food, whose chemical composition is show in Table 3, was used for analysis and measurements were performed in triplicate. Evaluation of inclusion percentage was determined at the beginning and at the end of technological operations and after a 12-month storage of the finished product, vacuum packed.

**Table 2.**
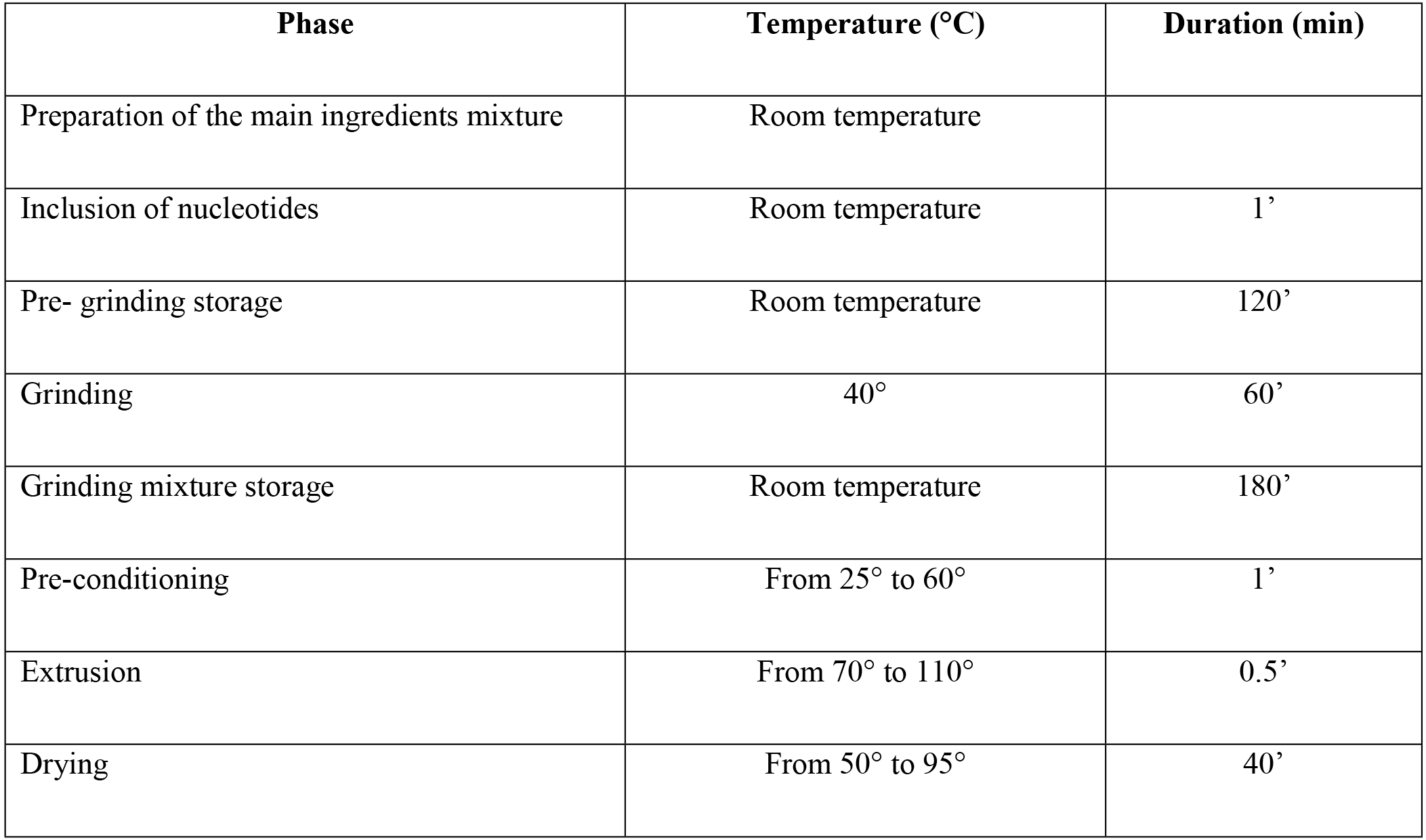
Operation preformed on dry food during phase and relative temperature and duration time.

| **Phase** | **Temperature (°C)** | **Duration (min)** |
| --- | --- | --- |
| Preparation of the main ingredients mixture | Room temperature |  |
| Inclusion of nucleotides | Room temperature | 1' |
| Pre- grinding storage | Room temperature | 120' |
| Grinding | 40° | 60' |
| Grinding mixture storage | Room temperature | 180' |
| Pre-conditioning | From 25° to 60° | 1' |
| Extrusion | From 70° to 110° | 0.5' |
| Drying | From 50° to 95° | 40' |

**Table 3.**
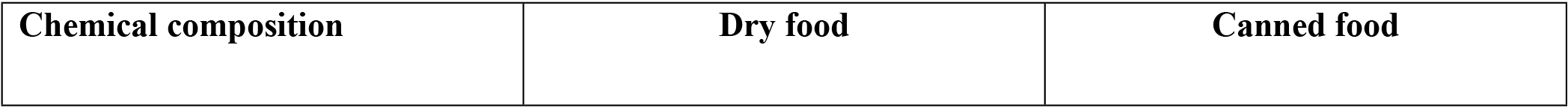
Chemical composition of dry and canned diets (g/100 g DM, except where specified)

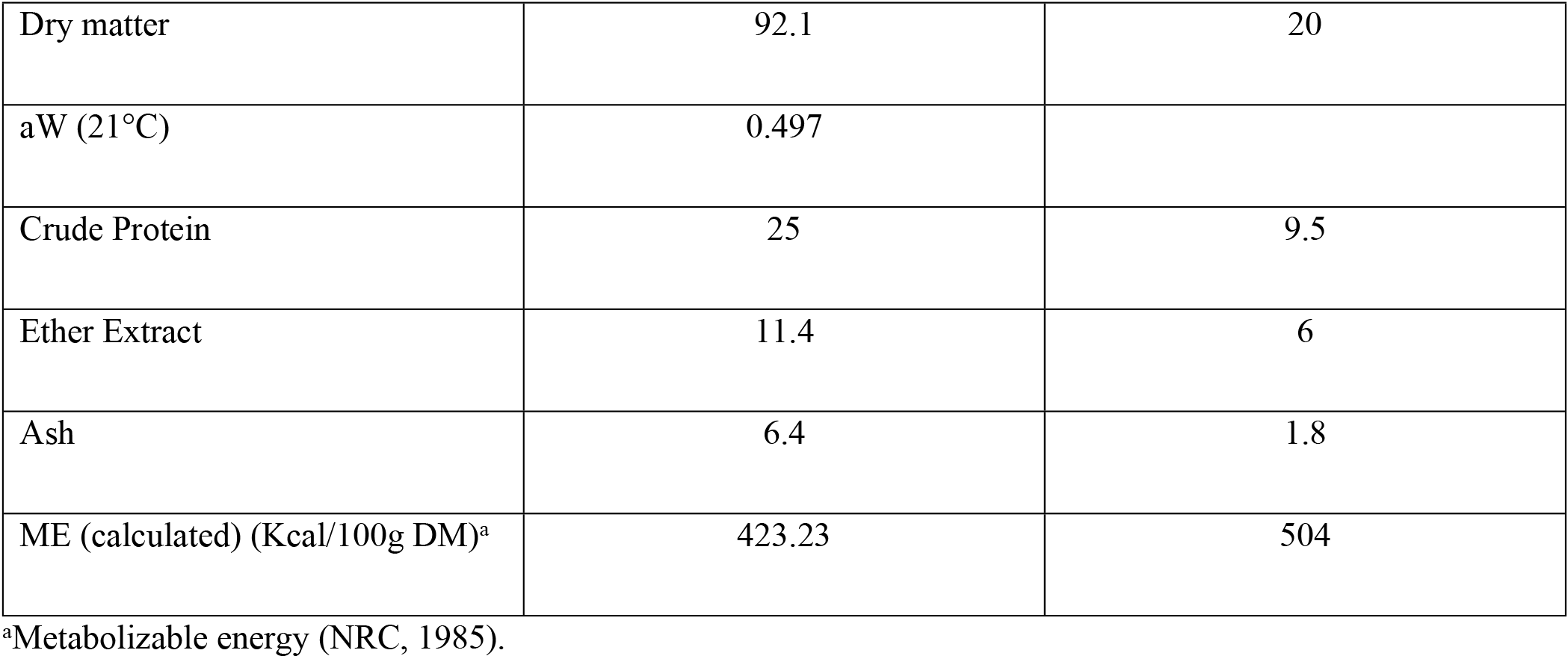

### Preparation of canned food

A commercial cat-pate formulation, whose chemical composition is show in Table 3, was prepared with a Prosol petMOD ™ percentage of inclusion of 0.4 g/100 on raw materials. Table 4 recapitulates the production flow occurred during dry food preparation. Product was packed in a 100 g aluminium tray, based on the evidence that this form is the one usually subjected to the most intense sterilization treatment. The nucleotides were inserted immediately before packaging to minimize their permanence in the unsterilized batch and to avoid any decay due to fermentation of the nucleotides by the bacterial flora present before sterilization, performed considering the unwanted food processing contaminants (FPCs) value, as suggested by Sevenich et al. [16]. The FPCs value, by convention, expresses the number of minutes necessary to obtain a lethal effect against a pathogen of reference at a temperature of 121.1 ° C, leading to sterilization. Evaluation of inclusion percentage was determined at the same time-points and under the same atmospheric conditions considered for dry food. Measurements were performed in triplicate.

**Table 4.**
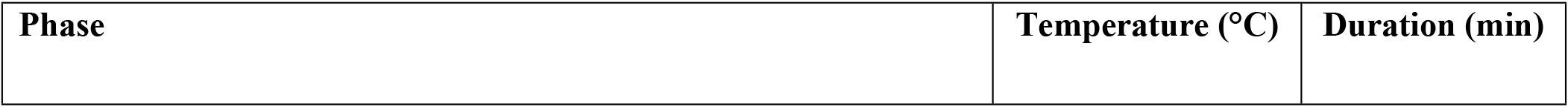
Operation preformed on canned food during phase 1 and relative temperature and duration time.

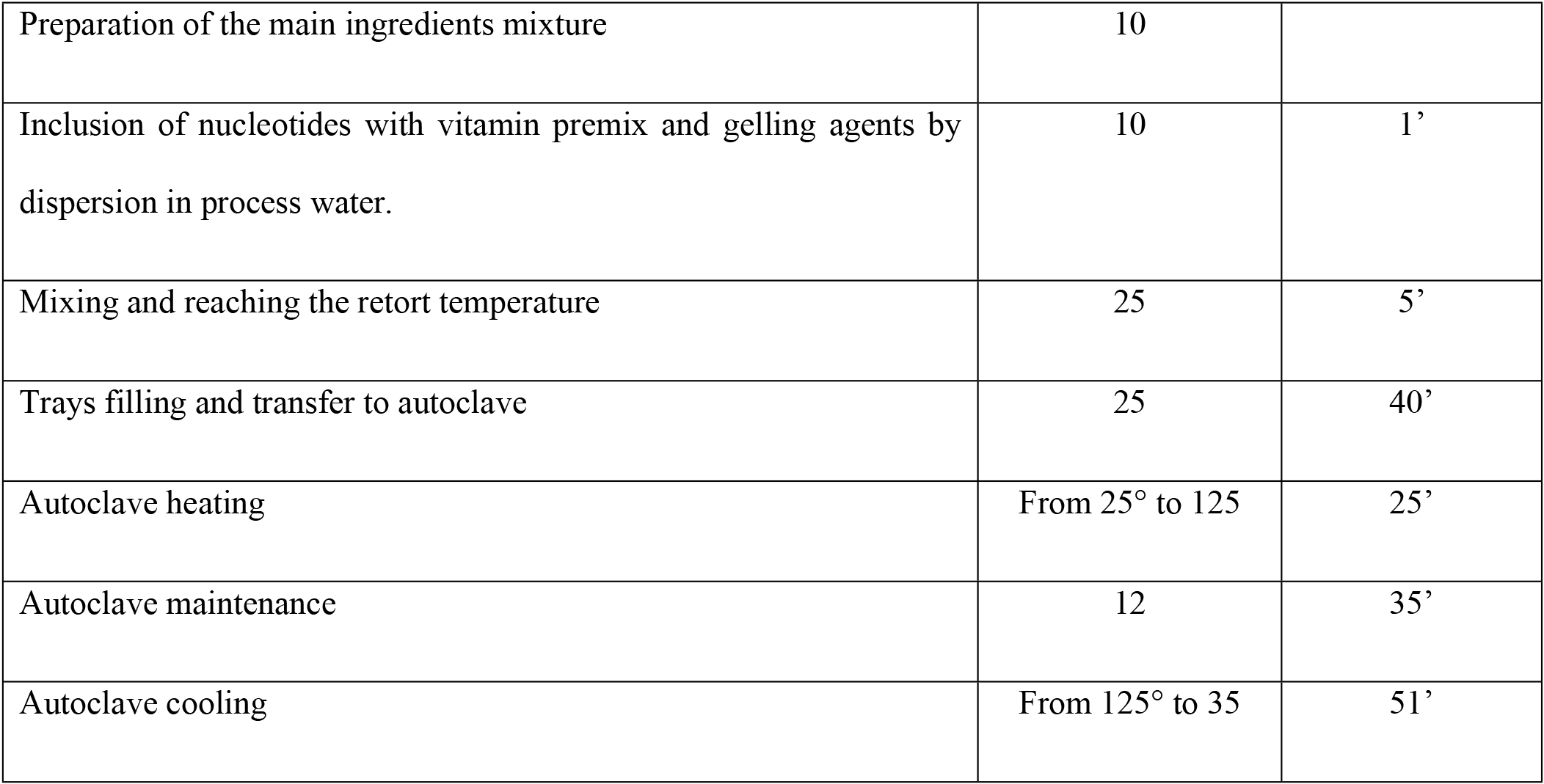

### Method for Nucleotides determination

At the beginning of the analyses 50 mg of each standard sample (AMP free acid, CMP free acid, GMP Na2, IMP Na2 and UMP Na) were weighted and diluted in HPLC water in order to reach a final volume of 100 ml. After a good solubilization, 10 ml of this solution were diluted in 50 ml of the same solvent. A volume of 10 µl of the final solution so obtained was injected in the HPLC system.

For dry food samples, 50 g of kibbles were weighted and grinded with a pestle and a mortar. A precise quantity of 1 g of grinded kibbles was weighted and dissolved in 100 ml of HPLC water; stir for 15 minutes and filtered on a paper filter of 30 microns. A filtrated volume of 10 µl was injected in the HPLC system.

For moist food, 50 g of pate were weighted, crushed with few drops of water and dissolved in 100 ml of HPLC water. The emulsion was settled for 30 min and 2 ml of supernatant were finally collected for HPLC injection.

The final report showing the different peaks and the related areas was printed at the end of the analyses. The retention time of the standards and of the sample were compared. The coefficient of variation (CV) was calculated and data with a CV ≤ 5% were validated.

Furthermore, the fraction of nucleotides coming from raw materials included in a dry and canned food characterized by the same analytic composition were measured during previous experiments (data not shown). These quantities were then deducted from values obtained at each time-point.

## Statistics

Statistical analysis was performed using SPSS, version 17.0 (SPSS Inc., Chicago, IL, USA). Nucleotides inclusion data were processed using the repeated measure model of ANOVA with Bonferroni post hoc test. A significant threshold was set at p<0.05. Results are reported as mean ± standard error.

## Results and Discussion

Nowadays, a growing trend in nucleotides supplementation is occurring in pets with the final goal of enhancing immune function and gut health, especially in young and debilitated animals. In these subjects, indeed, the cellular turnover is maximized and this aspect leads to an increased demand in dietary nucleotides as well. Segarra et al. [12] demonstrated that nucleotides improve biological markers of immune response in dogs and helped to a better-health status. This is particularly true for puppies since all tissues are in rapid growth and moreover the immune system is still incomplete [10–11]. The best way to achieve a dietary integration of nucleotides is to add them directly to a commercial food, even if the main source of these elements for healthy subjects is usually represented by meat and its by-product, which represent one of the main components of their maintenance diet. Despite the beneficial function of nucleotides administration is well recognized, little is still known about the capability of these elements to resist to technological processes. The present study was conducted to improve the knowledge in this field.

Recovery percentage of nucleotides in dry food at the end of technological process was 75% and 74% after a 12-month shelf-life, while in canned food it was 43% and 41%, respectively. Data are shown in Table 5. The gap existing between recovery percentages in dry and moist food are due to the fact that, during the sterilization process to which canned food is subjected, temperatures reached were much higher with a consequent greater deterioration of all products included [16]. Furthermore, recovery percentage of nucleotides could also be related to FPCs: higher it is and longer the duration of the thermal sterilization will be. FPCs we estimated for sterilization of our customized moist food was 69, which represent a standard value for canned pet food. It should however be said that the intensity of the treatment varies according to the volume of the container and the resistance opposed by its contents to the crossing by the heat. Pâtá was chosen precisely because, unlike chunks in sauce, it offers a greater resistance to heat penetration and therefore requires a greater use of thermal energy to obtain the same sterilizing effect. However, despite the amount of nucleotides degraded, their concentration after a 12 month shelf-life remained stable.

**Table 5.** Result of nucleotides determination during technological production and after a 12-month shelf-life. Data are given as mean ± standard deviation. (p < 0.05) DMB, dry matter bases.

|  | Beginning process | End product | 12-month shelf-life | *p value* |
| --- | --- | --- | --- | --- |
| **Dry food (g/100g DMB)** | 0.14 ± 0.01 | 0.12 ± 0.01 | 0.12 ± 0.01 | 0.005 |
| **Canned food (g/100g DMB)** | 0.82 ± 0.020 | 0.31 ± 0.07 | 0.29 ± 0.07 | 0.002 |

DMB, dry matter bases.

The technological process for dry food wasn’t able to significantly degrade raw materials leading to a higher recovery percentage as compared to moist food and a stable concentration during the 12-months shelf-life. This finding could be explained considering that during food production temperatures reached at the product core were lower than in canned food.

## Conclusions

In conclusion, nucleotides are raw materials authorized by Reg 01491-EN for inclusion in pet food that demonstrate different resistance to production process in dry or canned food. Pet food industries have to increase their awareness about this aspect, in order to ensure sufficient levels of nucleotides in the finished product able to exert beneficial effects on immune function and gastrointestinal system. Further studies are still warranted to optimize recovery percentage after inclusion in canned food and to evaluate their stability after a longer shelf-life period (i.e. 24 and 48 months).

## Author contributions

LP and DM designed the work. NR wrote the manuscript together with AC. NR also collected data and performed statistical analysis. LP and AC and DM performed a critical revision of the article. All authors gave final approval of the version to be published.

